# Changes in the gut microbiota and fermentation products associated with enhanced longevity in acarbose-treated mice

**DOI:** 10.1101/311456

**Authors:** Byron J Smith, Richard A Miller, Aaron C Ericsson, David C Harrison, Randy Strong, Thomas M Schmidt

## Abstract

**Background:** Treatment with the *α*-glucosidase inhibitor acarbose increases median lifespan by approximately 20% in male mice and 5% in females. This longevity extension differs from dietary restriction based on a number of features, including the relatively small effects on weight and the sex-specificity of the lifespan effect. By inhibiting host digestion, acarbose increases the flux of starch to the lower digestive system, resulting in changes to the gut microbiota and their fermentation products. Given the documented health benefits of short-chain fatty acids (SCFAs), the dominant products of starch fermentation by gut bacteria, this secondary effect of acarbose could contribute to increased longevity in mice. To explore this hypothesis, we compared the fecal microbiome of mice treated with acarbose to control mice at three independent study sites.

**Results:** Microbial communities and the concentrations of SCFAs in the feces of mice treated with acarbose were notably different from those of control mice. At all three study sites, the bloom of a single bacterial taxon was the most obvious response to acarbose treatment. The blooming populations were classified to the largely uncultured *Bacteroidales* family *Muribaculaceae* and were the same taxonomic unit at two of the three sites. Total SCFA concentrations in feces were increased in treated mice, with increased butyrate and propionate in particular. Across all samples, *Muribaculaceae* abundance was strongly correlated with propionate and community composition was an important predictor of SCFA concentrations. Cox proportional hazards regression showed that the fecal concentrations of acetate, butyrate, and propionate were, together, predictive of mouse longevity even while controlling for sex, site, and acarbose.

**Conclusion:** We have demonstrated a correlation between fecal SCFAs and lifespan in mice, suggesting a role of the gut microbiota in the longevity-enhancing properties of acarbose. Treatment modulated the taxonomic composition and fermentation products of the gut microbiome, while the site-dependence of the microbiota illustrates the challenges facing reproducibility and interpretation in microbiome studies. These results motivate future studies exploring manipulation of the gut microbial community and its fermentation products for increased longevity, and to test a causal role of SCFAs in the observed effects of acarbose.

## Background

The Interventions Testing Program (ITP) is a long-running, well-powered study of longevity enhancing interventions in genetically heterogeneous mice with identical protocols replicated at each of three study sites [1]. The drug acarbose (ACA) has been reproducibly shown in that study to increase mouse median lifespan with a larger effect in males than females [2, 3]. The largest increase was found when treatment began at 4 months, 22% in males and 5% in females [2], but the beneficial effect was still detectable in mice receiving ACA starting at 16 months [3]. The 90th percentile lifespan, a surrogate for maximum lifespan also shows benefits of ACA, with similar magnitudes in both male and female mice [2]. ACA is a competitive inhibitor of *α*-glucosidase and *α*-amylase, resulting in delayed intestinal breakdown of starch when taken with food and reduced postprandial increases in blood glucose. For these reasons, ACA is prescribed for the treatment of type 2 diabetes mellitus [4], and has also been shown to reduce the risk of cardiovascular disease in that population [5].

It is unclear whether the pathways by which ACA extends longevity in mice overlap with those affected by dietary restriction, but several observations have suggested critical differences [2]. Weight loss in ACA mice was more dramatic in females than in males, while the longevity effect is much stronger in males. By contrast, dietary restriction effects both weight and lifespan similarly in both sexes. Likewise, the response of fasting hormone FGF21 to ACA treatment was opposite in direction from that induced by dietary restriction. In female mice alone, ACA blocked age-related changes in spontaneous physical activity, while dietary restriction leads to dramatic increases in activity in both sexes. It is therefore justified to suspect that the effects of ACA on longevity are due to pathways distinct from dietary restriction.

Besides reducing the absorption of glucose from starch, inhibition of host enzymes by ACA results in increased flow of polysaccharide substrate to the lower digestive system [6]. ACA has been shown to raise the concentration of starch in stool [7, 6], and the observed increased excretion of hydrogen in breath [8, 9, 10, 11, 12] demonstrates that at least some of this substrate is fermented by the gut microbiota. The major byproducts of polysaccharide fermentation in the gut are hydrogen, CO_2_, and short-chain fatty acids (SCFAs), in particular acetate, butyrate, and propionate. Unsurprisingly, ACA treatment has been observed to increase acetate concentrations in human feces [13] and serum [14], as well as concentrations in portal blood and total amounts in rodent cecal contents [6]. Likewise, in some studies, ACA increased butyrate concentrations in human feces [7, 12, 15] and serum [14]. ACA also increased propionate concentrations in rat portal blood and total amount in cecal contents [6], as well as total output in feces in humans [13]. These changes were presumably due to changes in the activity and composition of microbial fermenters in the lower gut. Indeed, ACA was found to modulate the composition of the fecal bacterial community in prediabetic humans [16], and both increase the SCFA production potential inferred from metagenomes and lower fecal pH [17].

Impacts of ACA on microbial fermentation products are of particular interest because SCFAs produced in the gut are known to affect host physiology, with a variety of health effects associated with butyrate, propionate, and acetate (reviewed in [18] and [19]). Although butyrate and propionate are primarily consumed by the gut epithelium and liver, respectively, they are nonetheless detectable in peripheral blood, and acetate can circulate at substantially higher concentrations [20]. Four G-protein coupled receptors have been shown to respond to SCFAs with varying levels of specificity: FFAR2, FFAR3, HCA2, and OLFR78. All except OLFR78 are expressed in colonic epithelial cells, and each is expressed in a variety of other tissues throughout the body. Similarly, both butyrate and propionate act as histone deacetylase inhibitors which could have broad effects on gene expression through modulation of chromatin structure. In total, these pathways contribute to regulation of cellular proliferation, inflammation, and energy homeostasis, among other processes. The effects of ACA on fermentation products in the gut may, therefore, modulate its overall effects on host physiology.

Despite theoretical expectations and suggestive empirical results in other animal models, no study has looked for direct evidence that some of the longevity enhancing effects of ACA in mice are mediated by the gut microbiota and the SCFAs produced during fermentation. Here we test four predictions of that hypothesis: (1) ACA reproducibly modulates bacterial community composition; (2) the concentrations of SCFAs are increased in ACA-treated mice; (3) community structure is correlated with SCFAs and other metabolites in both control and treated mice; and (4) SCFA concentrations are predictive of lifespan. Fecal samples were analyzed from control and ACA treated mice enrolled in the ITP protocol at three, independent study sites.

## Results

### Study population

Sampled mice are representative of an underlying study population that recapitulates the previously observed sex-specific longevity effects of ACA. Across all three sites, ACA increased the median male survival of the underlying study population by 17% from 830 to 975 days (log-rank test *P* < 0.001). Female median survival increased 5% from 889 to 931 days (*P* = 0.003). These results are consistent with the increased longevity due to ACA previously reported [2]. Fecal samples from 48 mice at each of three study sites—The Jackson Laboratory (TJL), the University of Michigan (UM), and the University of Texas Health Science Center at San Antonio (UT)—were collected between 762 and 973 days of age with a balanced factorial design over sex and treatment group. Visual inspection of the overall survival curves (Fig. 1) confirmed that the longevity of mice sampled for microbiome analyses at UM and UT was representative of the other, surviving, unsampled mice. Samples from TJL were not matched to individual mice and longevity measures are not available for the subset described here.

**Figure 1.**
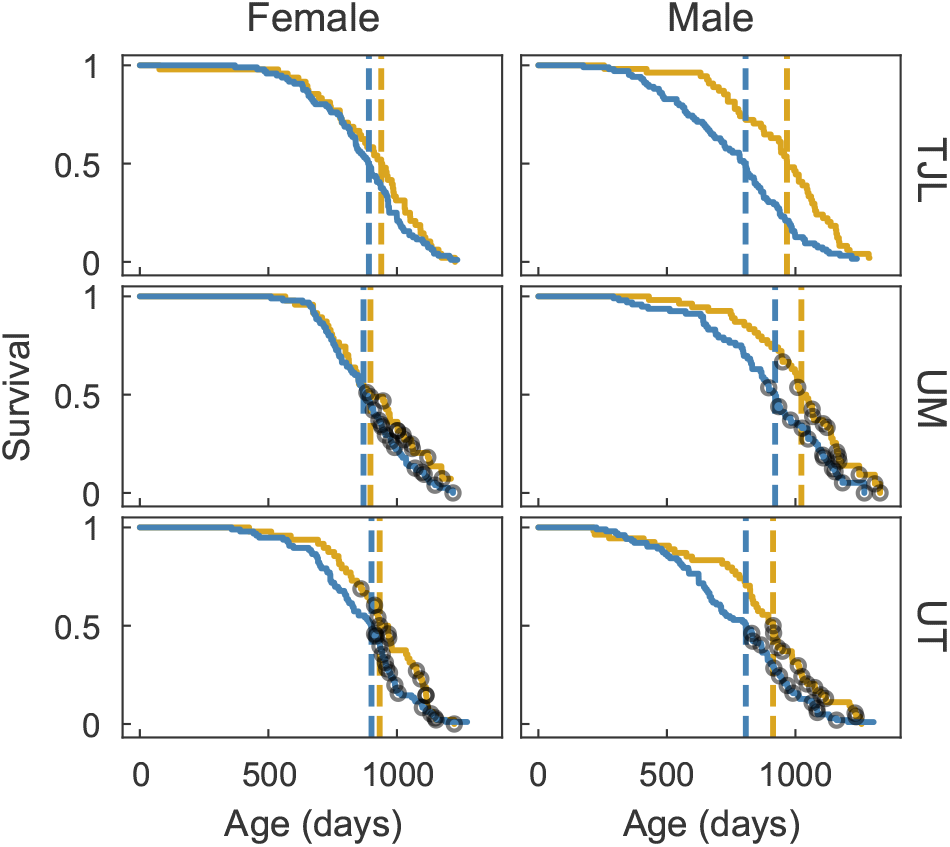
Survival curves of sampled cohort. Survival curves for mice fed a control diet (blue lines) or mice fed the same diet containing ACA (gold) at each of three sites: TJL, UM, and UT. Median longevity for each group of mice is indicated by a dashed vertical line. Black circles indicate the age at death for each of the sampled mice at UM and UT.

### Differences in fecal community in ACA-treated mice

ACA-treated mice had a substantially different microbial community composition from control mice at all three study sites. In a multivariate analysis of variance on site, sex, and treatment using Bray-Curtis dissimilarities and including all two-way interactions, significant effects were found for treatment (partial *r*^2^ = 9.6%, PERMANOVA *P* < 0.001) and site (partial *r*^2^ = 16.4%, *P* < 0.001), as well as their interaction (partial *r*^2^ = 3.4%, *P* < 0.001). These statistical results reflect the separation apparent in a principal coordinates ordination (see Fig. 2). A much smaller but still significant effect of sex (partial *r*^2^ = 1.0%, *P* = 0.014) was also identified, but there was no interaction between sex and treatment (*P* = 0.344). Despite the unbalanced design, significance of the PERMANOVA was not affected by changing the order of predictors. Based on a test of multivariate homogeneity of variances, dispersion differed between sites (PERMDISP *P* < 0.001) and sexes (*P* = 0.023), which may bias the PERMANOVA results, but did not differ between treatments (*P* = 0.425). The small effect of sex and the lack of significant interaction effects with treatment suggest that community composition itself, at the level of overall diversity, is not directly responsible for differential effects of ACA on longevity in male and female mice. However, the substantial differences in community composition due to treatment, while not surprising, suggests that the effects of ACA on the microbiome have the potential to modulate host health.

**Figure 2.**
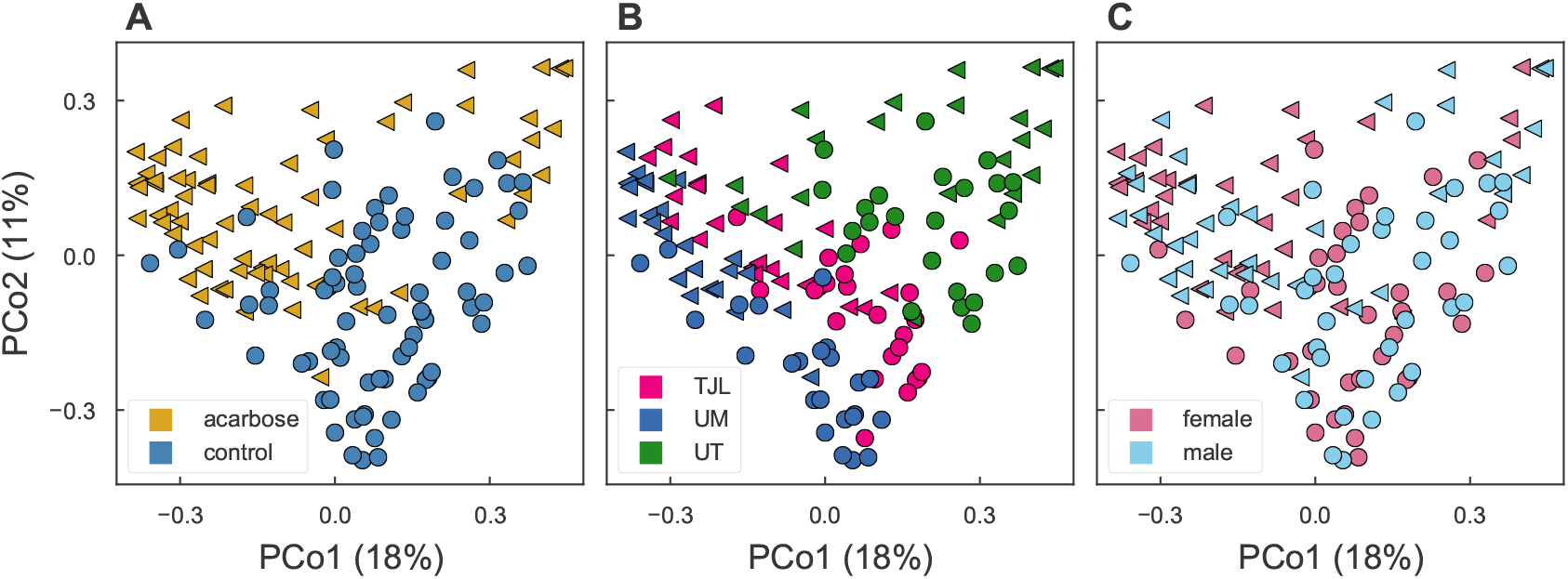
Microbial community composition. Fecal bacterial community composition in mice fed the control diet (circles) or the same diet supplemented with ACA (triangles). The two dominant principal coordinates, based on Bray-Curtis dissimilarities among community profiles, are plotted, and percent of variation explained by each is indicated in parentheses on the axes. The location of points in each panel is identical. Markers denote whether mice were treated (triangles) or controls (circles). In (A) points are colored by treatment: control mice (blue) and ACA-treated (gold), in (B) points are colored by site: TJL (pink), UM (blue), and UT (green), and in (C) points are colored by sex: male (light blue) and female (pink).

The fecal bacterial community in control mice was dominated by a handful of bacterial families (see Tbl. 1 for details). Across control mice at all three sites, a median of 30% of sequences were classified as members of the largely uncultured family *Muribaculaceae*—historically called the S24-7—belonging to the phylum Bacteroidetes. Other abundant families included the Lachnospiraceae (27%), Ruminococcaceae (14%), Lactobacillaceae (9%), and Erysipelotrichaceae (1%), all of which are classified in the phylum Firmicutes. More than 99.99% of sequences across all mice were classified at or below the family level. More detailed classification of sequences is provided in an additional file (see Appendix A).

At a 97% sequence similarity cutoff, 271 operational taxonomic units (OTUs) had a mean relative abundance across all samples of greater than 0.01% and an incidence of greater than 5%. Of these, the relative abundance of 113 OTUs differed between treated and control mice, correcting for a false discovery rate (FDR) of 5%. Together, these OTUs account for a median relative abundance of 54% across both control and treated mice. OTUs differing between sexes or reflecting an interaction between sex and treatment were a substantially less abundant. 5 OTUs were identified after FDR correction that differed significantly in relative abundance between male and female mice, accounting for a median, summed relative abundance of 6%. 7 OTUs were found to be subject to an interaction between treatment and sex, with a median relative abundances of 2%. Details of OTUs responsive to treatment, sex, and a treatment-by-sex interaction have been included in an additional file (see Appendix A).

Differences between control and ACA mice at TJL and UM were dominated by the increased abundance of a single OTU, OTU-1, which had a median relative abundance of 7.7% in control mice compared to 28.8% in ACA mice at TJL (Mann-Whitney U test *P* < 0.001), and 10.4% compared to 39.0% at UM (*P* < 0.001) (see Fig. 3). At UT, OTU-1 was higher in ACA-treated mice—a median of 5.4% and 11.0% in control and treated mice, respectively—but this increase was not statistically significant (*P* = 0.344). Instead, a different OTU, designated OTU-4, was strongly affected by ACA treatment at UT, with a median relative abundance of 6.3% in control mice that increased to 25.6% in ACA-treated mice (*P* = 0.007). OTU-4 was nearly absent at TJL and UM, with only one mouse out of 95 having a relative abundance above 0.1%, compared to 39 out of 48 mice at UT. Differences in abundance between sexes were not observed for OTU-1 at TJL or UM, but at UT results were suggestive of an increased abundance of OTU-1 in females (*P* = 0.076) and an increased abundance of OTU-4 in males (*P* = 0.060). Interestingly, their combined abundance did not differ between males and females (*P* = 0.344) at UT. Both OTU-1 and OTU-4 were classified as members of the *Muribaculaceae,* and subsequent phylogenetic analysis confirmed this placement (Appendix C). OTU-1 and OTU-4 are approximately 90% identical to each other and to the most closely related cultivar (DSM-28989) over the V4 hypervariable region of the 16S rRNA gene. These OTUs are notable both for their high abundance overall, as well as the large difference between control and ACA-treated mice. It is surprising that OTU-4 is common and differentially abundant at UT, while remaining rare at both of the other sites, suggesting that local community composition modulates the effects of ACA. While OTU-1 is made up of multiple unique sequences, the composition within the OTU does not differ substantially with ACA treatment (see Fig. 4).

**Figure 3.**
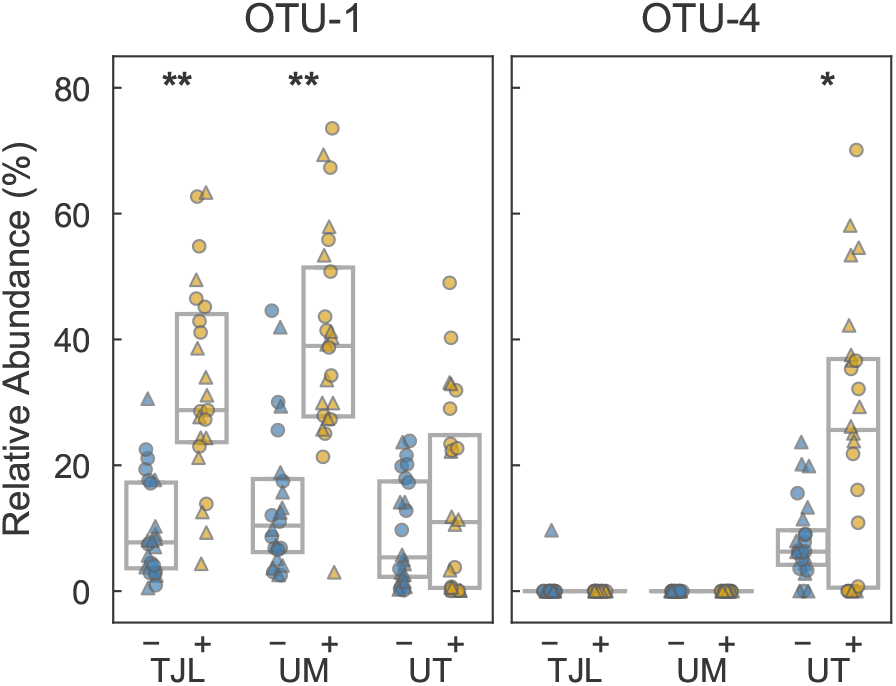
Abundance of dominant OTUs. Relative abundance of the 16S rRNA gene from two OTUs abundant in ACA-treated mice. Points in each panel correspond with samples collected from individual mice at each of three replicate study sites. Samples were obtained from mice fed either the control diet (blue) or the same diet supplemented with ACA (gold). Markers indicate the sex of the mouse: male (triangle) or female (circle). Boxes span the interquartile range and the internal line indicates the median. (*: *P* < 0.05, **: *P* < 0.001 by Mann-Whitney U test).

**Figure 4.**
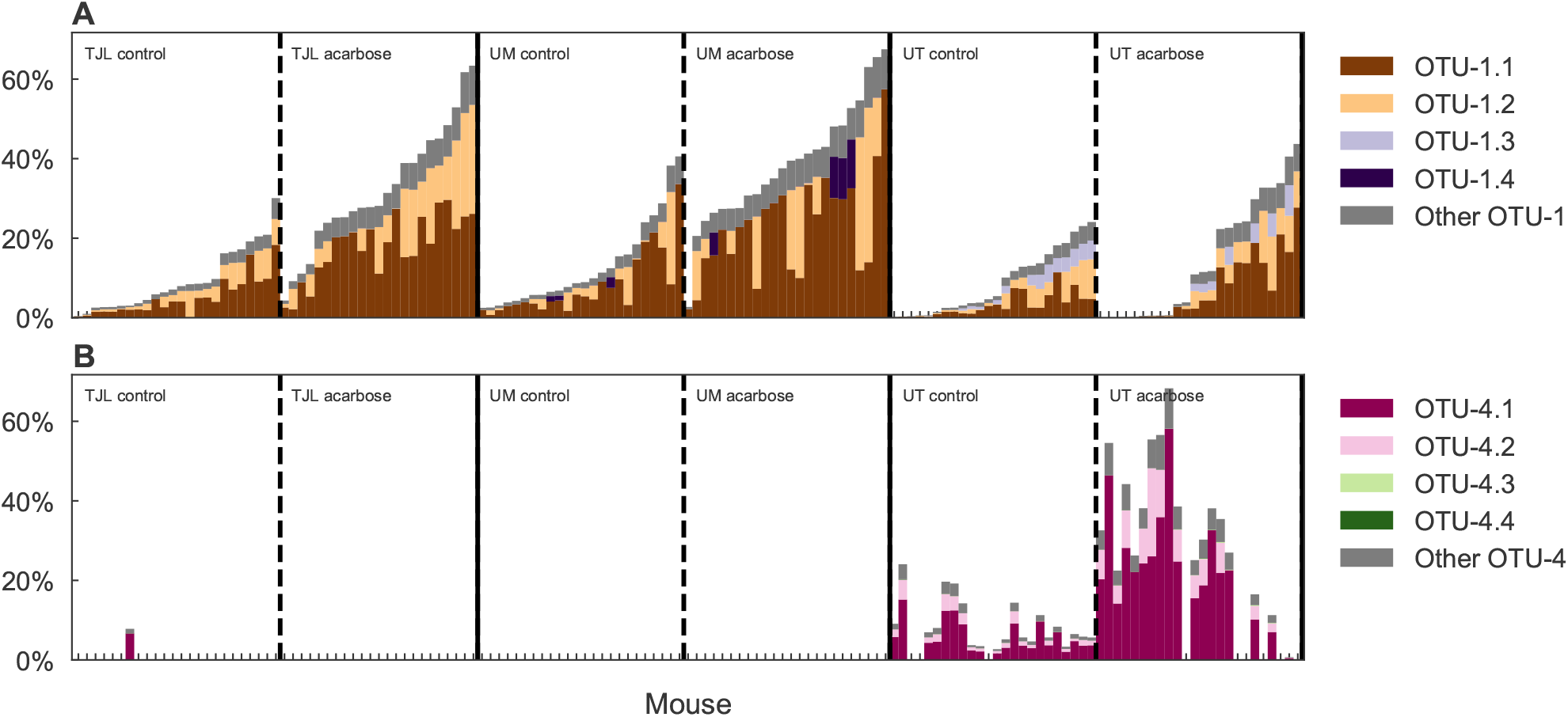
Relative abundance across samples of unique sequences. Distribution of common unique sequences clustered into (A) OTU-1 and (B) OTU-4. Colors are assigned to the top four most common unique sequences within each OTU and all remaining sequences from that OTU are assigned the color gray. Stacked bars in each position represent individual mice sampled for this study, and reflect the relative abundance of unique sequences in that sample. Mice are grouped into sites and then treatments, and finally by the total abundance of OTU-1.

The increased relative abundance of OTU-1 and OTU-4 in mice treated with ACA appears to be due to greater abundance of these sequences, and is not explained solely by a decrease in other groups. The abundance of taxa in control and treated mice was compared based on the recovery of spiked-in standard relative to the sequence of interest. The median combined spike-adjusted abundance of 16S rRNA gene copies from OTU-1 and OTU-4 was 4.3 times greater per gram of feces in ACA-treated mice compared to controls (Mann-Whitney U test *P* < 0.001), suggesting a corresponding increase in population density.

While OTU-1 and OTU-4 are classified to the same family and are similarly affected by ACA, other OTUs in the *Muribaculaceae* have decreased abundance in treated mice. The combined relative abundance of all other OTUs in the family— excluding OTU-1 and OTU-4—was 8.3% in treated mice versus 16.8% of sequences in control mice (Mann-Whitney U test *P* < 0.001). The median combined spike-adjusted abundance of all other *Muribaculaceae* OTUs was 0.5 times the median in control mice (*P* = 0.001), suggesting a decrease in the population density of these taxa. This is consistent with competition between OTUs in this family.

Three of the five most abundant families all exhibit decreased relative abundance in ACA treated mice (see Tbl. 1). However, the large increase in abundance of OTU-1 and OTU-4 suggests that some changes in the relative abundance of other taxa may be the result of compositional effects, rather than decreased density. For instance, although the relative abundance of Ruminococcaceae was lower in ACA-treated mice, the spike-adjusted abundance was little changed (*P* = 0.327), emphasizing the value of this complementary analysis. Conversely, decreased relative abundance was matched by decreased spike-adjusted abundance for both the Lactobacillaceae (*P* = 0.014) and the Erysipelotrichaceae (*P* = 0.063).

**Table 1.** Abundance of common bacterial families.

| family | % control<sup>a</sup> | % ACA<sup>a</sup> | ACA : control<sup>b</sup> |
| --- | --- | --- | --- |
| *Muribaculaceae* | 30.4 (21.5, 43.3) | 48.1<sup>**</sup> (35.3, 61.8) | 1.8<sup>**</sup> |
| *Lachnospiraceae* | 26.6 (16.3, 41.6) | 23.9<sup>†</sup> (9.6, 37.4) | 1.3 |
| *Ruminococcaceae* | 14.2 (9.0, 19.0) | 11.6<sup>*</sup> (6.9, 15.6) | 1.1 |
| *Lactobacillaceae* | 9.5 (1.2, 17.0) | 2.6<sup>*</sup> (1.0, 8.2) | 0.31<sup>*</sup> |
| *Erysipelotrichaceae* | 1.4 (0.3, 6.2) | 0.5<sup>*</sup> (0.2, 2.2) | 0.42<sup>†</sup> |

<sup>a</sup> Median and interquartile range of the relative abundance of the top five most abundant bacterial families in control and ACA-treated mice
<sup>b</sup> the ratio of median spike-adjusted abundances in ACA-treated mice versus control mice
<sup>†</sup> $P < 0.1$ via Mann Whitney U test
<sup>\*</sup> $P < 0.05$
<sup>\*\*</sup> $P < 0.001$

ACA-treated mice exhibited decreased fecal community diversity. The median Chao1 richness estimate was decreased from 229 in control mice to 199 in treated mice (Mann-Whitney U test *P* < 0.001). The Simpson’s evenness index was also lower in ACA mice: 0.044 versus 0.075 in controls (*P* < 0.001). This reduced richness and evenness is not surprising given the much greater abundance of a single OTU in treated mice at each site. To understand changes in diversity while controlling for compositional effects, we measured the effect of ACA ignoring counts for OTU-1 and OTU-4. While Simpson’s evenness was not decreased by treatment in this fraction of the community (*P* = 0.26), the Chao1 richness—subsampling to equal counts *after* partitioning—was (*P* = 0.005), suggesting that the bloom of OTU-1 and OTU-4 may have, in fact, resulted in the local extinction of rare community members.

### Changes in fecal metabolite concentrations

Long-term ACA treatment affects metabolite profiles, increasing concentrations of the SCFAs in feces (see Fig. 5). Butyrate concentrations were increased from a median of 3.0 mmols/kg wet weight in control mice to 4.9 in ACA-treated mice (Mann-Whitney U test *P* < 0.001). Propionate concentrations were also increased: a median of 1.1 in controls compared to 2.3 with ACA (*P* < 0.001). Median acetate concentrations were higher, 16.2 mmols/kg versus 12.9 in controls, but a Mann-Whitney U test did not surpass the traditional p-value threshold (*P* = 0.073). The summed concentrations of acetate, butyrate, and propionate was greater in ACA-treated mice, with a median concentration of 25.4 mmols/kg versus 19.0 mmols/kg in control mice (*P* = 0.003). This confirms our predictions given the expectation of greater availability of polysaccharide substrate for fermentation. Indeed, median glucose concentration was also increased from 5.3 to 10.3 (*P* < 0.001). Concentrations of formate, valerate, isobutyrate, and isovalerate were generally below the detection limit. Fresh pellet weight was increased from a median of 36 to 74 mg (*P* < 0.001) Fecal starch content was not measured, but pellets from ACA-treated mice had a noticeably chalky appearance.

**Figure 5.**
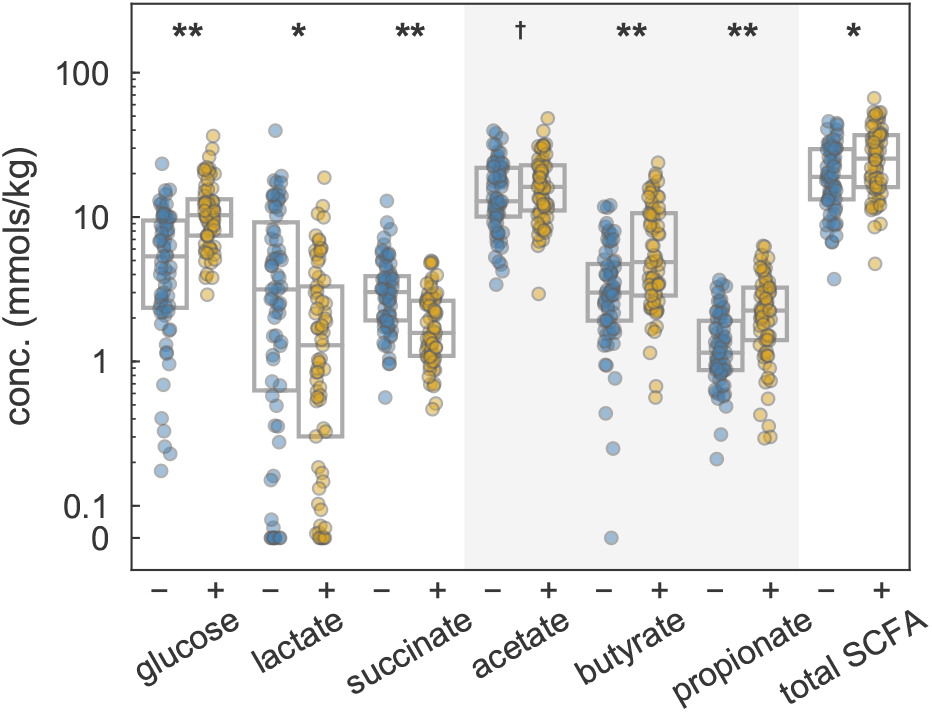
Fecal metabolites. Concentrations of metabolites in feces obtained from mice fed either the control diet (blue) or the same diet supplemented with ACA (gold). Boxes span the interquartile range and the internal line indicates the median. The shaded region highlights the three major SCFAs produced by microbial fermentation of polysaccharides in the gut, and the sum of their concentrations is plotted as “total SCFA”. Above 0.1 mmols/g, concentrations are plotted logarithmically, (†: *P* < 0.1, *: *P* < 0.05, **: *P* < 0.001 by Mann-Whitney U test).

Butyrate as a molar percentage of total SCFA was modestly greater in the ACA mice, a median of 19% in control mice was increased to 22% in treated mice (Mann-Whitney U test *P* < 0.001), as was propionate: 7% in control, 10% in treated (*P* = 0.006), while acetate was decreased from 73% in controls to 66% in treated mice (*P* < 0.001).

In contrast to the three measured SCFAs and glucose, both succinate and lactate concentrations were decreased. Median lactate was decreased from 3.2 mmols/kg in control mice to 1.3 in ACA-treated mice (*P* = 0.003), and succinate from 3.0 to 1.6 (*P* < 0.001). It is surprising that these fermentation intermediates are reduced, given the expected increase in available polysaccharide. It is possible that their concentrations reflect greater consumption in downstream pathways, or perhaps ACA directly inhibits the metabolism and growth of relevant bacteria; such effects have been previously reported for *in vitro* fermentations of starch with human fecal slurries [7].

Differences in SCFAs between sexes are particularly interesting given the greater longevity effects of ACA in male mice. For propionate, a sex-by-treatment interaction was found (ANOVA *P* = 0.023), but butyrate and acetate had no such effect. This interaction results in a larger difference in propionate concentrations for male mice (from 1.4 mmols/kg in control to 2.7 in ACA) than for female mice (from 1.0 to 1.9) with ACA treatment. The significance of the interaction term was not corrected for multiple testing, and therefore additional studies would greatly increase our confidence in this result. Fecal metabolite concentrations also varied over study site and sex of the mouse; effects of ACA stratified by site and sex are visualized in Fig. 7.

### Community predictors of fecal SCFA concentrations

Community composition was correlated with metabolite concentrations in both control and ACA-treated mice. Numerous strong correlations were detected between the spike-adjusted abundance of 16S rRNA copies from the most common bacterial families and the concentrations of SCFAs and lactate. Notably, *Muribaculaceae* abundance was particularly strongly correlated with propionate concentrations in both control (Spearman’s *ρ* = 0.36, *P* = 0.002; see Fig. 6) and ACA mice (*ρ* = 0.64, *P* < 0.001). Likewise, Lachnospiraceae were correlated with butyrate (*ρ* = 0.61 in control and 0.77 in ACA, *P* < 0.001 for both), and Lactobacillaceae with lactate concentrations (*ρ* = 0.63 in control and 0.67 in ACA, *P* < 0.001 for both). Strikingly, concentrations of acetate and butyrate were especially correlated with each other (*ρ* = 0.67 in control and 0.80 in ACA, *P* < 0.001 for both). Although our study was not an unambiguous test, these results support the hypothesis that the fecal metabolite response to treatment is dependent on the population density of relevant microbes in the gut community. Similarly, environmental and host factors that promote or inhibit the growth of particular community members would be expected to modulate the response.

**Figure 6.**
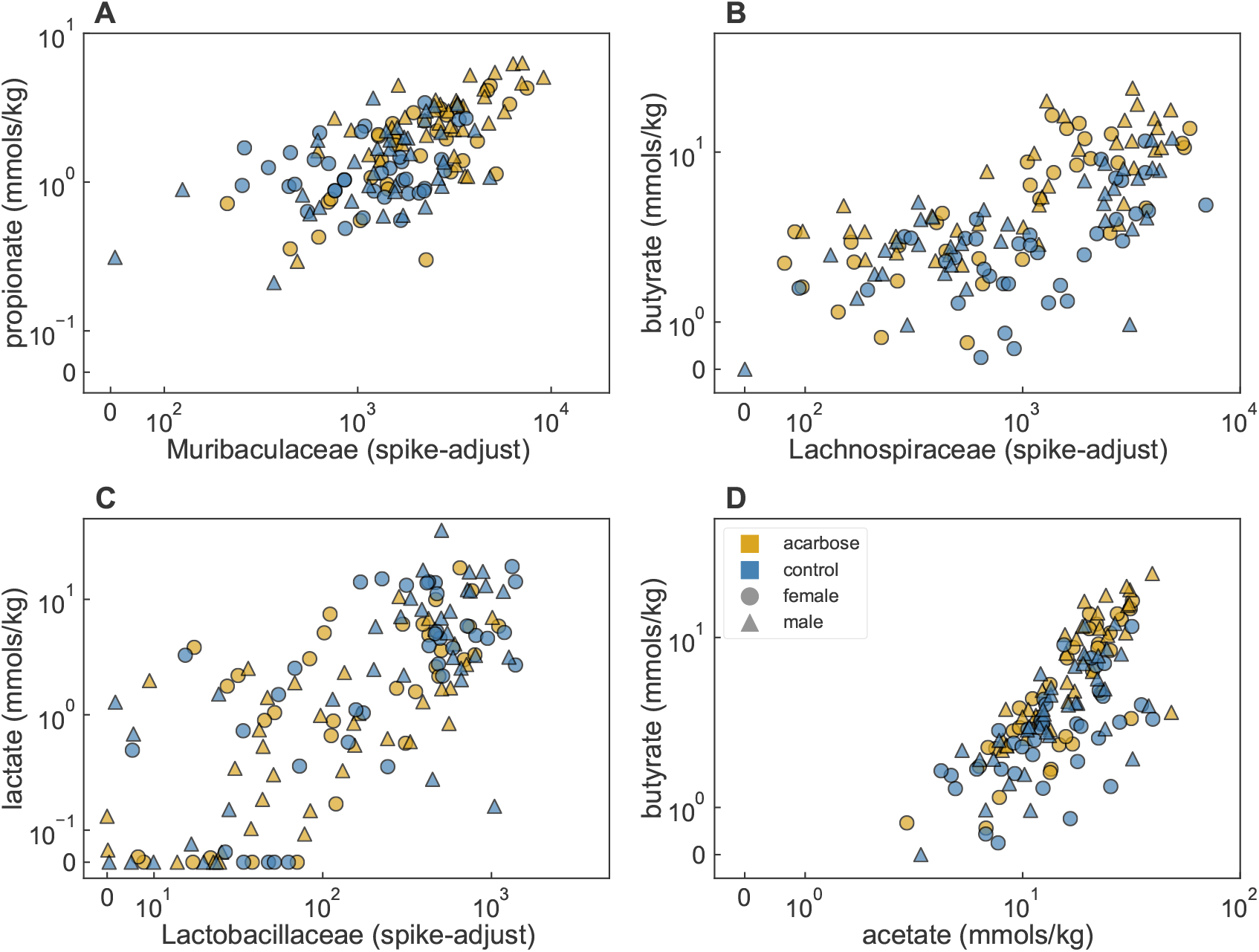
Microbiota/metabolite correlations. Correlations among metabolite concentrations in feces and family level, spike-adjusted 16S rRNA gene abundances. Points correspond with samples collected from individual mice and colors indicate whether they were obtained from mice fed the control diet (blue) or the same diet supplemented with ACA (gold). Markers indicate the sex of the mouse: male (triangle) or female (circle). Metabolite concentrations are reported normalized to feces wet weight, and abundances are in spike-equivalent units. Values are on a linear scale between 0 and the subsequent tick label, above which, points are plotted logarithmically.

To identify key players in these associations, we examined the relationship between metabolite concentrations and the spike-adjusted abundances of OTUs. Based on a LASSO regression, the abundances of a number of OTUs can be used to predict concentrations of propionate, butyrate, acetate, and lactate even after accounting for treatment, sex, and study site. Consistent with the correlations found between *Muribaculaceae* abundance and propionate, OTU-1 and OTU-4 were identified as predictors of increased propionate, along with a third taxon, OTU-5, also classified as a member of the family. For both butyrate and acetate, OTUs classified as members of both the Lachnospiraceae and Ruminococcaceae were most predictive of increased concentrations. Unsurprisingly, the most abundant OTU classified as a member of the Lactobacillaceae, OTU-2, was found to be highly predictive of increased lactate. However, 8 OTUs were also associated with decreased lactate concentrations, most of which were among those associated with increased butyrate and acetate. Among other explanations, this is consistent with these taxa either being inhibited by lactate or being lactate utilizers, which are likely to be producing SCFAs as secondary fermentation products. Regression coefficients for each metabolite are included in an additional file (see Appendix A). Overall, results were both consistent with *a priori* expectations, and useful for generating hypotheses about which taxa might be associated with the generation of fermentation products.

### Fecal SCFA concentrations as predictors of longevity

Given the documented health benefits of SCFAs in the gut (reviewed in [18]) and their increased levels in ACA-treated mice, we tested the relationship between the acetate, butyrate, and propionate concentrations in feces, and the lifespan of individual mice. Lifespans of fecal donors were not available for mice at TJL, so survival analyses were carried out only with UM and UT mice, and effect sizes are reported for SCFAs as standardized hazard ratios (HRs). Due to the reduced number of mice sampled for this study, data were pooled across sexes and sites. The shared effects of the design parameters—treatment, sex, and study site—on both SCFAs and longevity, were accounted for by including terms for these covariates as well as their two and three-way interactions. Analyses reinforcing our interpretations are discussed in Appendix D. Tested individually against this null model, an association between longevity and propionate was found (standardized HR of 0.727, *P* = 0.031), but no relationship was found with butyrate (*P* = 0.240) or acetate (*P* = 0.742). However, when the model was fit with all three SCFAs simultaneously, each was found to be associated with longevity (*P* = 0.012, 0.030, 0.042 for propionate, butyrate, and acetate, respectively). Coefficients for SCFA covariates in this full model suggest a positive association with longevity for both propionate and butyrate (standardized HR of 0.674 and 0.586, respectively). Interestingly, a negative association was found with acetate (standardized HR of 1.576) using this model. The discrepancy between this result and the lack of association when butyrate and acetate are each tested alone likely reflects the strong positive correlation between acetate and butyrate concentrations, masking their individual, opposing associations with longevity. The overall fit of the full model was improved compared to the null model with only design covariates (likelihood ratio test, *P* = 0.023).

## Discussion

ACA, by inhibiting the enzymes responsible for starch degradation in the small intestine, is expected to increase the availability of this polysaccharide to the microbiome. The resulting increase in SCFA production may contribute to the effects of ACA on health. Despite previous observations in humans and rats that ACA results in substantial changes to the community structure [16, 7] and fermentation products [6, 7, 12, 15] of the gut microbiota, a link between these effects on the microbiome and longevity has not been established. Here we present the first study to combine bacterial community surveys with measurement of fecal metabolites in ACA treated mice, as well as the first to pair these data with lifespan, allowing us to explore the role of the microbiome in increased longevity.

Our results confirm all four predictions that we set out to test: ACA was found to affect both (1) the composition of bacterial communities and (2) SCFAs in mouse feces, (3) the abundances of individual taxonomic groups were associated with concentrations of fermentation products, and (4) the concentrations of fecal SCFAs were associated with variation in mouse longevity.

While it is unsurprising that an increased flux of starch to the large intestine affected the gut microbiota and their fermentation products, some changes were especially pronounced. The increased relative abundance in ACA-treated mice of the dominant OTU—OTU-1 at UM and TJL and OTU-4 at UT—was dramatic: one or the other was increased approximately 4-fold at all three sites and in multiple samples more than half of sequences belonged to these OTUs. A cursory BLAST search reveals that sequences identical to OTU-1 have been previously recovered in published studies (e.g. [21] and [22]); in [23] the sequence was found at high relative abundance in the brains of mice that had undergone sepsis. It was notable that OTU-1 did not respond to ACA at UT, while its increased abundance was so striking at UM and TJL. Our results appear to constrain the potential explanations for this observation. OTU-1 was present and abundant at all three sites; The abundance in control mice was lowest at UT, although the median there was still greater than 5%. While it is not possible with the data presented here to rule out genomic differences of OTU-1 among sites, a similar composition of unique 16S rRNA gene sequences made up this cluster at all three. On the other hand, OTU-4 was at very low abundance, with no reads in a majority of samples, at UM and TJL where OTU-1 did respond to ACA. These results suggest that both OTUs respond to ACA in the same way, with OTU-4, when it is sufficiently abundant, inhibiting the response of OTU-1, potentially through resource competition. Both OTUs are in the same family, the *Muribaculaceae*, but are not the same species or genus by the traditional similarity thresholds, sharing only 90% identity over the sequenced fragment. The differential response of these OTUs among sites illustrates the importance of each site’s local “metacommunity” in determining the microbial community’s response to environmental perturbations.

Pronounced differences in the resident microbial communities of different hosts may contribute to challenges in translating results from mice and other model organisms to humans. A comparison of bacterial community composition in feces in prediabetic people before and during a 4-week ACA treatment period did not reveal changes of the magnitude reported here [16], although this may reflect the limited duration of treatment. Interestingly, in that study *Lactobacillaceae* abundance increased with ACA, while we observed this family to be depleted in treated mice. The abundance of the *Muribaculaceae* was not reported. Although this family is common in mice and have been previously shown to respond to diet, the clade is substantially less abundant in most human samples [24]. However, the prevalence of *Muribaculaceae* may be under-reported in the literature, as the Ribosomal Database Project [25] does not include the family and classification using this database assigns sequences to the *Porphyromonadaceae* instead [26]. Historically, two other names have also been used for this clade: the “S24-7” (from an early environmental clone [27]), and “ *Candidatus Homeothermaceae”* (proposed in [24]). While the isolation of one *Muribaculaceae* cultivar has recently been published, *Muribaculum intestinale* YL27 [28], and as of this writing several draft genomes are available for unpublished isolates (e.g. see whole genome shotgun sequencing projects NWBJ00000000.1 and NZ_NFIX00000000.1), additional cultivars will be vital for understanding the function and ecology of the family. Nonetheless, genomes assembled from metagenomes suggest that populations of *Muribaculaceae* are equipped with fermentation pathways to produce succinate, acetate, and propionate, and that the family is composed of metabolic guilds, each specializing on the degradation of particular types of polysaccharides: plant glycans, host glycans, and *α*-glucans [24]. This suggests that the *Muribaculaceae* may occupy a similar set of niches in mice as do *Bacteroides* species in humans. The *Muribaculaceae* and *Bacteroides* are both in the order *Bacteroidales*. *Bacteroides* also specialize in the fermentation of polysaccharides [29], and at least some of the most common species in the human gut are known to produce succinate, acetate, and propionate from the fermentation of polysaccharides [30, 31, 32, 18]. Unlike the patterns observed here for *Muribaculaceae* in mice, the abundance of *Bacteroides* decreased with ACA treatment in one study in humans [16], suggesting that the microbially mediated effects of ACA may fundamentally differ between these hosts.

Besides hypotheses based on genome content, the correlation between total *Muribaculaceae* abundance and propionate concentrations and the specific association with OTU-1 and OTU-4 found in the LASSO analysis suggest that both OTUs, and perhaps other *Muribaculaceae* species in this study, ferment starch to propionate. This also supports the hypothesis, discussed above, that both OTUs occupy overlapping niches. Although increased butyrate concentrations have been frequently reported with ACA treatment [7, 14, 6, 12, 15], elevated concentrations of propionate have been observed in just one previous study using portal blood in rats [6]. Studies in humans have instead found decreased or no change in fecal [13, 7, 12] or serum [14] propionate concentrations with ACA. Decreased propionate has been attributed to preferential production of butyrate from starch fermentation [33, 34] or inhibition of propiogenic bacteria by ACA [7]. Our observation of increased propionate was robust and reproduced at all three sites. If this conflicting result reflects both the greater initial abundance and enrichment in our study of the *Muribaculaceae*—especially OTU-1 and OTU-4— it demonstrates the value of measuring both community composition and metabolite concentrations in the same samples.

SCFAs are commonly suggested to act as intermediaries between the gut microbiota and host physiology [18]. While our study was not designed to provide a causal test of effects of SCFAs on longevity, and the power of our analysis was limited, a statistical association between SCFA concentrations and mouse lifespan supports an interpretation that is consistent with an extensive literature on the health benefits of butyrate and propionate [18]. In addition, that SCFA concentrations were associated with longevity above and beyond the effects of ACA, study site, and sex, further supports this hypothesis. It is somewhat surprising, however, that a single fecal sample taken, in some cases, several months before death, could be predictive of longevity. The association reported here could reflect other, unmeasured, changes in the gut microbiome or host physiology. Concentrations of metabolites in feces are an integration of both production and consumption rates along the length of the lower gut, and may not reflect host exposure nor the strength of host physiological response. It is also important to note that, since all mice in this study at the time of sampling were of an age close to the median lifespan of control individuals, the results are only relevant to mechanisms of aging in late-life and should not be extrapolated to young mice. Experimental tests of a causal role for SCFAs in longevity will be challenging, as they likely require controlled manipulation of intestinal SCFAs for the lifetime of a mouse.

Due to the preferential enhancement of longevity by ACA in male mice we sought to identify aspects of the gut microbiome that responded differently in male and female mice (i.e. interaction effects), as these might suggest mechanistic explanations for differences in longevity effects [2]. While we do not believe that the magnitude of sex-by-treatment interactions observed for various aspects of the microbiome were sufficiently pronounced to fully explain the differential effect of ACA on lifespan, our search was limited by sample size, variability between study sites, and the large number of features being tested. Nonetheless, ACA was found to increase propionate concentrations more in male mice than females, a statistically significant pattern before correction for multiple testing, and the relative abundance of a handful of OTUs seem to have also been subject to an interaction.

Mechanisms unrelated to the gut microbiome have been proposed for the effects of ACA on lifespan. Because ACA reduces the postprandial glucose spike observed in mice and humans, hypotheses emphasizing the reduction of harmful effects associated with these transient surges have been most commonly invoked. Studies of UM-HET3 mice given ACA from 4 to 9 months of age suggested that mean daily blood glucose levels are minimally affected, but that absorption of glucose was both slower and longer lasting [2]. Interestingly, fasting insulin level in ACA-treated males are much lower than those in control males, consistent with an improvement in insulin sensitivity [2]. This reduction was not seen in females, where insulin levels in controls were lower than in control males and similar to those in ACA-treated males [35, 2], presenting one possible explanation for the stronger longevity benefit in males. Still, the connection between this modulation of postprandial glucose— with or without improved insulin sensitivity—and extended longevity is still far from certain.

The work presented here explores a different hypothesis: that health benefits in ACA mice are related to changes in the activity of microbial communities in the gut associated with the increased influx of starch, and possibly attributable to known health effects of microbial metabolites, including SCFAs. The changes described here in both community composition and fermentation products due to ACA treatment, along with the statistical association between fecal SCFA concentrations and longevity, are consistent with this hypothesis, and provide a stepping-stone for future studies. Interestingly, SCFAs themselves have well documented effects on glucose homeostasis (reviewed in [36] and [37]). The two explanations are therefore not mutually exclusive, and the effects of ACA on longevity may be mediated by both glucose physiology and microbial activity in the gut.

## Conclusions

Here we have tested four predictions of a proposed model connecting ACA to lifespan via the gut microbiome. We demonstrate that ACA reproducibly modulates the composition of the microbiota, as well as the concentrations of fermentation products, increasing the abundance of butyrate and propionate. In addition, we provide evidence that the structure of the microbial community is an important factor in the composition of metabolites produced. Finally, we show an association between SCFA concentrations in feces and survival, suggesting a role of the microbiome in the life-extending properties of ACA. Together, these results encourage a new focus on managing the gut microbiota for host health and longevity.

## Methods

### Mouse housing and ACA treatment

Mice were bred and housed, and lifespan was assessed as described in [2]. Briefly, at each of the three study sites, genetically heterogeneous, UM-HET3, mice were produced by a four-way cross as previously described in [38]. After weaning, mice were fed LabDiet^®^ (TestDiet Inc.) 5LG6 produced in common batches for all sites. At 42 days of age, electronic ID chips were surgically implanted and treatment randomly assigned to each cage housing four mice to a cage for females and three to a cage for males. ACA-treated mice were fed the same chow amended with 1000 ppm ACA (Spectrum Chemical Manufacturing Corporation) from 8 months of age onwards. Mice were transferred every 14 days to fresh, ventilated cages with water provided in bottles. Colonies at all three sites were assessed for infectious agents four times each year, and all tests were negative for the entire duration of the study.

### Sample collection and processing

Fresh fecal pellets were collected directly from mice between 762 and 973 days of age and frozen at −80°C. We did not control the time of day at collection. While differences in age and collection time could have added variability to SCFA concentrations, both were similarly distributed for the different treatment groups, so they are unlikely to confounded our analyses. To eliminate potential cage effects from cohoused mice [39], samples were obtained from no more than one randomly selected mouse per cage. A total of 144 samples were collected from 12 male and 12 female mice in both control and ACA treatment groups at each of the three sites. Samples were shipped on dry ice and then stored, frozen, until processing. For approximately the first half of samples, we extracted the soluble fraction by homogenizing pellets with 200 μL of nuclease-free water. For the remaining samples, we instead used a 1:10 ratio (weight:volume), with a maximum volume of 1.5 ml. This was found to improve quantification in higher weight samples. While SCFA concentration estimates were higher when using the amended protocol, the order of sample extraction was fully randomized, so it is unlikely to have biased our interpretations. Homogenized samples were centrifuged at 10,000 × g for 10 minutes to separate soluble and solid fractions. The supernatant was then serially vacuum filtered, ultimately through a 0.22 μm filter, before HPLC analysis. The solid fraction was frozen prior to DNA extraction. Four samples were excluded from chemical analysis and one from DNA analysis due to technical irregularities during sample processing.

Prior to DNA extraction, fecal pellet solids were thawed and, where necessary, subsampled for separate analysis. To move beyond relative abundance, solids were weighed and spiked with 10 μL aliquots of prepared *Sphingopyxis alaskensis* strain RB2256—an organism not found in mouse feces—in order to compare 16S rRNA gene abundance between samples [40, 41]. The spike was prepared as follows: a 1:200 dilution of a stationary phase *S. alaskensis* culture was grown at room temperature for approximately 44 hours in R2B medium with shaking. This culture was harvested at a final OD420 of 0.72 before being rinsed in PBS and resuspended—5-fold more concentrated—in 20% glycerol in PBS (v/v). Aliquots of these cells were stored at −20°C before extraction and sequencing. Spiked fecal samples were homogenized in nuclease free water at a ratio of 1:10 (w/v). DNA was extracted from 150 μL of this mixture using the MoBio PowerMag Microbiome kit.

### Chemical analysis

The chemical composition of samples was assessed on a Shimadzu HPLC (Shimadzu Scientific Instruments) equipped with an RID-10A refractive index detector. 30 μL injections were run on an Aminex HPX-87H column (Bio-Rad Laboratories, Hercules, CA) at 50°C with 0.01 N H_2_SO_4_ mobile phase and a flow rate of 0.6 ml/minute. External standards were run approximately daily containing acetate, butyrate, formate, glucose, lactate, propionate, and succinate at 8 concentrations between 0.1 mM to 20 mM. Due to the complexity of the chromatogram, the identity and area of retained peaks was curated manually, assisted by the LC Solutions Software (Shimadzu Scientific Instruments) Standard curves were fit using weighted regression (inverse square of the concentration), and, for all compounds except propionate, without an intercept.

### 16S rRNA gene sequencing and analysis

The V4 hypervariable region of the 16S rRNA gene was amplified from extracted DNA (as described in [42]), and sequenced on an Illumina MiSeq platform using a MiSeq Reagent Kit V2 500 cycles (cat# MS-102-2003). Amplicon sequences were processed with MOTHUR (version 1.39.4 [43]) using a protocol based on the 16S standard operating procedures [42]. Scripts to reproduce our analysis can be found at [44]. After fusing paired reads, quality trimming, and alignment to the SILVA reference database (Release 132 downloaded from [45]). The vast majority of 16S rRNA gene sequences were between 244 and 246 bp. Sequences were classified using the method of Wang et al. [46] as implemented in MOTHUR and with the SILVA non-redundant database as a reference [47]. We clustered sequences into OTUs using the OptiClust method [48] at a 97% similarity threshold. We counted and removed sequences classified as *S. alaskensis,* the spiked-in standard, before further analysis. We did not attempt to assess the exact number of 16S rRNA gene copies spiked into samples. Instead, spike-adjusted abundance was defined in units based on the standardized spike (μL spike equivalents / g sample) and estimated using the formula:

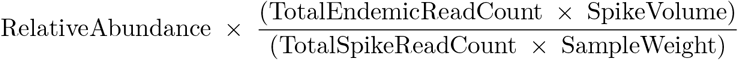

Family level abundance was calculated as the summed abundance of all sequences clustered in OTUs classified to that family. OTU counts were randomly subsampled to the minimum number of reads before calculating Chao1 richness, but were not subsampled for other analyses. A single, independent realization of random subsampling was used for each richness calculation. A search of the NCBI non-redundant nucleotide database for related sequences from cultured bacteria was carried out using the BLASTn web tool [49] searching the non-redundant nucleotide database with default parameters and excluding sequences from uncultured organisms.

### Statistical analysis

A 0.05 p-value threshold was used to define statistical significance, with values below 0.1 considered “suggestive”. Except where specified, p-values are not corrected for multiple testing. Due to the risk of violating distributional assumptions, univariate comparisons between groups were done using the non-parametric Mann-Whitney U test. Differences in multivariate community composition and dispersion were tested using PERMANOVA (adonis) and PERMDISP (`betadisper`) respectively, both implemented in the vegan package (version 2.4-6 [50]) for the R programming language. Bray-Curtis dissimilarity was used as the *β*-diversity index.

Differences in the relative abundance of individual OTUs were surveyed using the DESeq2 package (version 1.18.1 [51]) for R, and fitting a model that included terms for treatment, sex, site, and the interaction between treatment and sex. So as to keep valuable distributional information, all OTUs found in at least two samples were included in the initial analysis, with p-values calculated using a Wald test. However, FDR correction using the Benjamini-Hochberg procedure [52] excluded “rare” OTUs—those with mean relative abundance less than 0.01% or detected in fewer than 5% of samples—in order to maintain statistical power by independently reducing the number of tests.

Interactions between sex and treatment in fecal SCFAs were assessed for log-transformed concentrations. The small number of zeros were replaced with half the lowest detected concentration for that metabolite. Interactions were tested in an ANOVA that also included terms for site, sex, and treatment. LASSO regressions of the three SCFAs and lactate against spike-adjusted OTU abundances were performed using the scikit-learn library for Python (version 0.18.2 [53]) and log-transformed concentrations after adjustment for site, sex, and treatment. OTUs detected in more than 5% of samples and with mean abundance greater than 0.01% were included. The LASSO parameter was determined by randomized 10-fold crossvalidation, optimizing for out-of-bag R^2^. For each metabolite we confirmed that OTU abundance information improved predictions by testing the Spearman’s rank correlation between true values and out-of-bag predictions of the best model using a Student’s t-distribution approximation [54] and a *P* = 0.05 significance threshold. While this type of regularized regression is primarily useful for constructing predictive models, and biological interpretation can be challenging, non-zero regression coefficients are suggestive of covariates that are among the most strongly associated with a response. Proportional hazards regression was carried out using of the survival package (version 2.41-3 [55]) for R, and the day of fecal sampling as the entry time. All sampled mice were dead at the time of analysis and right-censoring was therefore not used. Standardized HRs reported for SCFAs are based on concentrations that have been centered around 0 and scaled to a standard deviation of 1.

## Appendix A: Description of Additional file - OTU details table

Summary statistics, associations with design parameters and metabolites, and inferred taxonomic identity of common bacterial OTUs. Table includes all OTUs with mean relative abundance greater than 0.01% and found in greater than 5% of samples. Associations with treatment, sex, and a treatment-by-sex interaction are included in columns labeled “treatment_effect”, “sex_effect”, and “interact_effect”, respectively. Columns labeled “propionate”, “butyrate”, and “acetate” include fitted coefficients from the respective LASSO regression. Formatting has been included to facilitate exploration, with background colors indicating the sign and magnitude of coefficients and colored circles indicating the statistical significance of results after FDR correction.

## Appendix B: Supplemental figure

**Figure 7.**
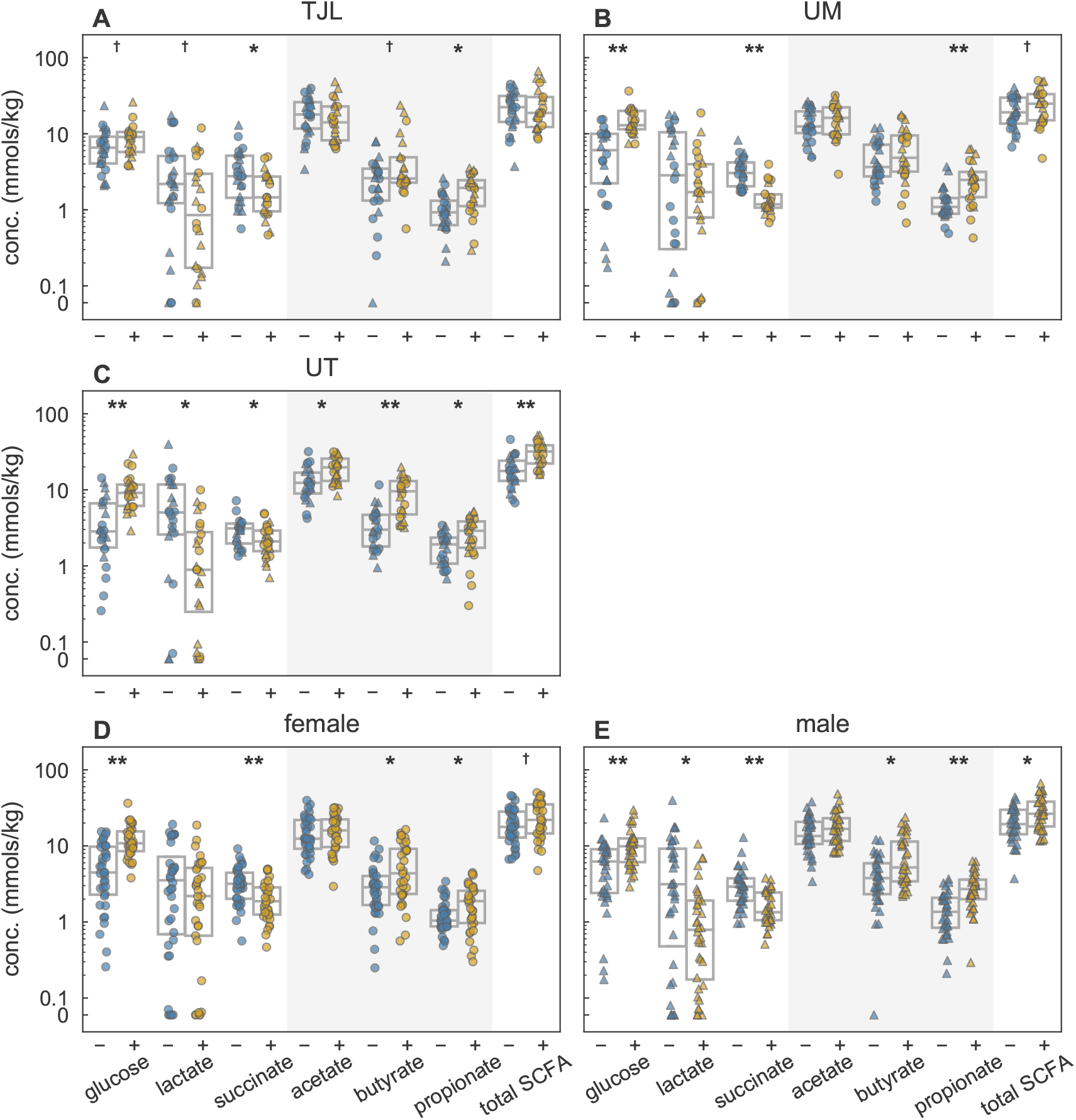
Concentrations of metabolites stratified by site and sex. Chemical analysis of feces obtained from mice fed either the control diet (blue) or the same diet supplemented with ACA (gold) within individual (**A-C**) study sites and (**D-E**) sexes. Markers indicate the sex of the mouse: male (triangle) or female (circle). Boxes span the interquartile range and the internal line indicates the median. The shaded region highlights the three major SCFAs produced by microbial fermentation of polysaccharides in the gut, and the sum of their concentrations is plotted as “total SCFA”. Above 0.1 mmols/g, concentrations are plotted logarithmically. Significance symbols reflect differences in concentration between treated and control mice (†: *P* < 0.1, *: *P* < 0.05, **: *P* < 0.001 by Mann-Whitney U test).

## Appendix C: Taxonomic analysis of two dominant OTUs

The most conspicuous difference in the gut microbiota induced by ACA was a site-specific effect in two populations of bacteria both classified as members of family *Muribaculaceae* (Fig. 3). OTUs were classified based on an approximately 240 bp fragment of the 16S rRNA gene in the V4 hypervariable region. Using this fragment, we applied several lines of evidence to confirm that OTU-1 and OTU-4 are both members of the *Muribaculaceae* and that they are genetically distinct from cultured relatives. Classification of sequences using the method of Wang et al. [46] and the SILVA non-redundant database as a reference [47], identified both OTU-1 and OTU-4 as members of the family with 100% bootstrap support. While use of the RDP training set [25] Version 14 instead assigned these sequences to the family Porphyromonadaceae this is presumably because the *Muribaculaceae* are not recognized as a taxon in the RDP (previously reported by [26]), nor are alternative names for the clade (“S24-7” or “*Homeothermaceae”).* A follow-up phylogenetic analysis of representative amplicon sequences from two dominant OTUs was carried out using approximate maximum likelihood estimation implemented in the FastTree software (version 2.1.8 [56]) using the generalized time reversible model with twenty discrete rate categories (`-gtr -gamma` options). Approximate maximum likelihood phylogenetic estimation, using a selection of type strains in the order Bacteroidales, places OTU-1 and OTU-4 in a clade with representatives of the *Muribaculaceae* with >95% support for the topology of that node (see Fig. 8). While such a short sequence fragment is unlikely to perfectly recapitulate phylogeny—indeed, tree topology was generally weakly supported and was sensitive to both the choice of reference sequences and the evolutionary model used—we are nonetheless satisfied with the evidence for assignment of both OTU sequences to this clade; besides exceptions in the Porphyromonadaceae, Marinilabiliaceae, and *Bacteroides,* our phylogenetic reconstruction largely matches a recently proposed taxonomy of the Bacteroidales [24].

OTU-1 and OTU-4 represent uncultured genera. Over the analyzed sequence they have 89% and 92% identity, respectively, to *Muribaculum intestinalis* strain YL27, the first cultured representative of the *Muribaculaceae* [28]. A BLAST search against the NCBI non-redundant nucleotide collection did not find higher sequence similarity to any other cultured bacteria. Representative sequences for OTU-1 and OTU-4 share nucleotides at only 22 out of 244 positions (91%).

**Figure 8.**
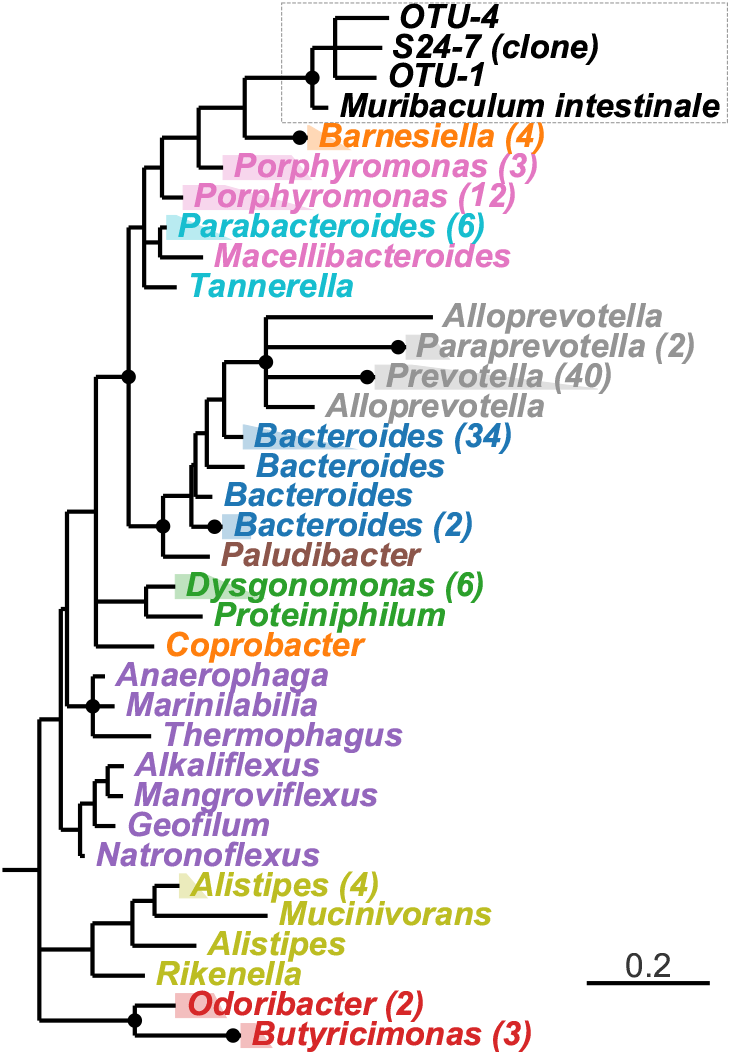
Phylogenetic characterization of OTU-1 and OTU-4. Estimated phylogeny based on approximately 240 bp of the 16S rRNA gene V4 hypervariable region and more than 130 type strain reference sequences spanning the diversity of order Bacteroidales. Branch lengths are in units of expected substitutions per site. The tree is rooted by a Flavobacteriales out-group (not shown). Reference taxa are labeled with genus designations according to the SILVA database. When multiple representatives from the same genus have been folded together, the number of sequences is reported in parentheses. Nodes with Shimodaira-Hasegawa local support over 95% are indicated with black circles and nodes with support less than 70% have been collapsed to polytomies. The dashed box encloses taxa inferred to be within the *Muribaculaceae.* The taxon labeled ‘S24-7 (clone)’ (GenBank: AJ400263.1) is the environmental sequence by which the clade was originally identified, and by which it was historically named [27], while *Muribaculum intestinalis* (GenBank: KR364784.1) is the first cultured representative [28]. Label colors indicate a recently proposed family membership of each reference: Prevotellaceae (gray), Barnesiellaceae (orange), Porphyromonadaceae (pink), Dysgonomonadaceae (green), Bacteroidaceae (blue), Tannerellaceae (light blue), Marnilabilaceae (purple), Marinifilaceae (red), Paludibacteraceae (brown), and Rikenellaceae (yellow) [24].

## Appendix D: Expanded survival analysis

Proportional hazards regression was used in this study to demonstrate an association between SCFA concentrations in feces and the longevity of mice. Given the limited number of samples for which matched chemical and survival data are available, statistical testing of associations with SCFAs were carried out in the pooled dataset. Therefore, to account for known effects of treatment, sex, and site, the primary null model used in this study includes terms for all main, two, and threeway interactions of the design covariates: treatment, sex, and study site. A priori, a number of these terms were expected to be non-zero, based on ITP findings from previous cohort years [2, 3]. Indeed, when survival data from all of control and ACA treated mice in this cohort were analyzed together, effects were detected that recapitulated these expectations, including: increased longevity of females, increased longevity with ACA treatment, and increased longevity of control males at UM (see Tbl. 2).

**Table 2.** Fitted coefficients for experimental design covariates in the full ITP cohort data.

| Term | log(HR) | Std. Error | P |
| --- | --- | --- | --- |
| ACA | -0.530 | 0.170 | 0.002 |
| Female | -0.266 | 0.143 | 0.063 |
| TJL | -0.024 | 0.144 | 0.865 |
| UM | -0.598 | 0.155 | 0.000 |
| ACA:Female | 0.230 | 0.245 | 0.349 |
| ACA:TJL | -0.225 | 0.242 | 0.353 |
| ACA:UM | 0.003 | 0.254 | 0.992 |
| Female:TJL | -0.017 | 0.204 | 0.934 |
| Female:UM | 0.580 | 0.213 | 0.006 |
| ACA:Female:TJL | 0.351 | 0.349 | 0.313 |
| ACA:Female:UM | -0.054 | 0.361 | 0.881 |

Interestingly, some—though not all—of these effects were still evident when analyzing the much smaller data set of mice from which we collected fecal samples.

**Table 3.**
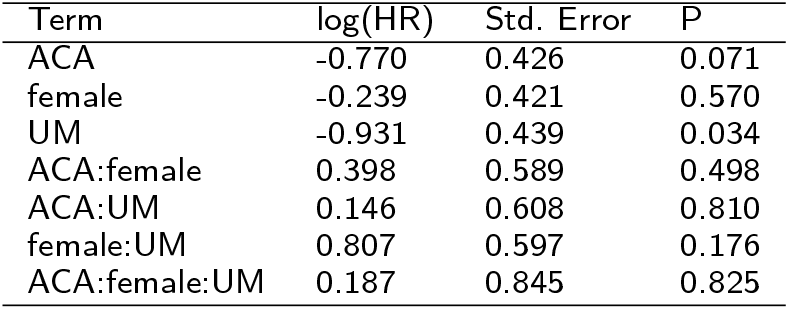
Fitted coefficients for experimental design covariates in the sampled mice.

| Term | log(HR) | Std. Error | P |
| --- | --- | --- | --- |
| ACA | -0.770 | 0.426 | 0.071 |
| female | -0.239 | 0.421 | 0.570 |
| UM | -0.931 | 0.439 | 0.034 |
| ACA:female | 0.398 | 0.589 | 0.498 |
| ACA:UM | 0.146 | 0.608 | 0.810 |
| female:UM | 0.807 | 0.597 | 0.176 |
| ACA:female:UM | 0.187 | 0.845 | 0.825 |

This increased our confidence that, despite the age of the mice at the time of entry and the limited sample size, the associations with SCFA concentrations reflect patterns that could be seen in the full cohort.

As described in the main results, adding terms for the concentrations of three SCFAs improved the fit of the model (see Tbl. 4).

**Table 4.** Fitted coefficients for experimental design covariates and SCFAs in the sampled mice.

| Term | log(HR) | Std. Error | P |
| --- | --- | --- | --- |
| propionate | -0.292 | 0.116 | 0.012 |
| butyrate | -0.119 | 0.055 | 0.030 |
| acetate | 0.062 | 0.030 | 0.042 |
| ACA | -0.124 | 0.488 | 0.799 |
| female | -0.476 | 0.433 | 0.272 |
| UM | -1.232 | 0.484 | 0.011 |
| ACA:female | 0.095 | 0.597 | 0.874 |
| ACA:UM | 0.241 | 0.654 | 0.713 |
| female:UM | 0.913 | 0.614 | 0.137 |
| ACA:female:UM | 0.111 | 0.850 | 0.896 |

Interestingly, the positive association betwen ACA and longevity is distinctly weakened when SCFAs are included (estimated coefficient went from −0.770 without SCFAs to −0.124 with). Although we do not carry out a formal path analysis, this is consistent with the causal effects of ACA on longevity being mediated by SCFA concentrations.

Survival analysis is potentially sensitivity to deviations from the assumptions of the Cox family of models [57]. We therefore tested proportionality and linearity assumptions relevant to our main findings. The Cox proportional hazards model assumes that the hazard associated with each covariate is proportional across the full set of ages at which mice are being tracked—e.g. that the proportional decrease in the risk of death for mice treated with ACA is equal from the first entry time to the last exit. We checked this assumption using a test of the correlation between the scaled Schoenfeld residuals and Kaplan-Meier transformed survival times (implemented as the `cox.zph` function in the survival package for R, [55]) and found no evidence for deviations for any of the design parameters nor for the included SCFAs.

**Table 5.** Tests of non-proportionality for experimental design covariates and SCFAs in the sampled mice.

| Term | Correlation | P |
| --- | --- | --- |
| propionate | -0.127 | 0.155 |
| butyrate | 0.039 | 0.633 |
| acetate | 0.012 | 0.885 |
| ACA | 0.047 | 0.630 |
| female | 0.065 | 0.518 |
| UM | -0.036 | 0.701 |
| ACA:female | -0.090 | 0.401 |
| ACA:UM | 0.044 | 0.643 |
| female:UM | -0.028 | 0.778 |
| ACA:female:UM | 0.037 | 0.726 |
| GLOBAL | — | 0.922 |

Similarly, visual inspection of residual plots did not provide any evidence of deviations from linearity assumptions inherent to the model (Fig. 9).

**Figure 9.**
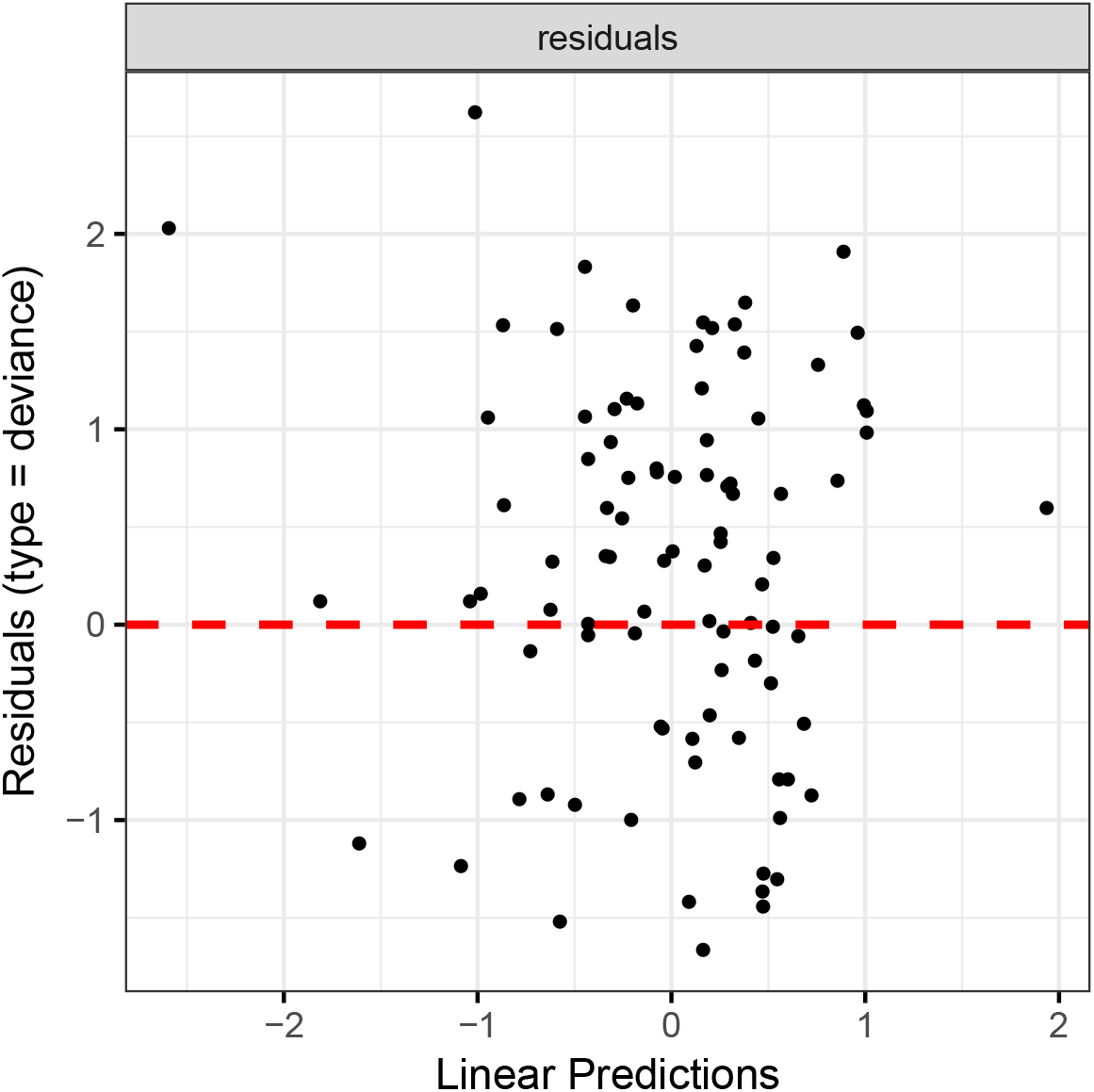
Proportional hazards model residuals. Deviance residuals versus predicted log-hazard in a model of mouse survival that includes all design parameters (site, sex, and treatment) as well as the three SCFAs: propionate, butyrate, and acetate.

While regression coefficients can be directly interpreted as a proportional increase or decrease in hazard of death at all time points, in the general case, this does not equate to a proportional change in expected survival time. It can therefore be challenging to understand the magnitude of the survival effect on expected lifespan. To provide a more intuitive demonstration of the size of the effect, we compared predicted survival curves for ACA treated, male mice at UM, with different SCFA concentrations characteristic of two existing individuals, based on a hypothetical scenario in which mice were alive at 830 days of age (i.e. a conditional expected survival curve; see Fig. 10). These simulated results demonstrate the strength of the association between SCFAs and survival over observed differences in concentrations.

**Figure 10.**
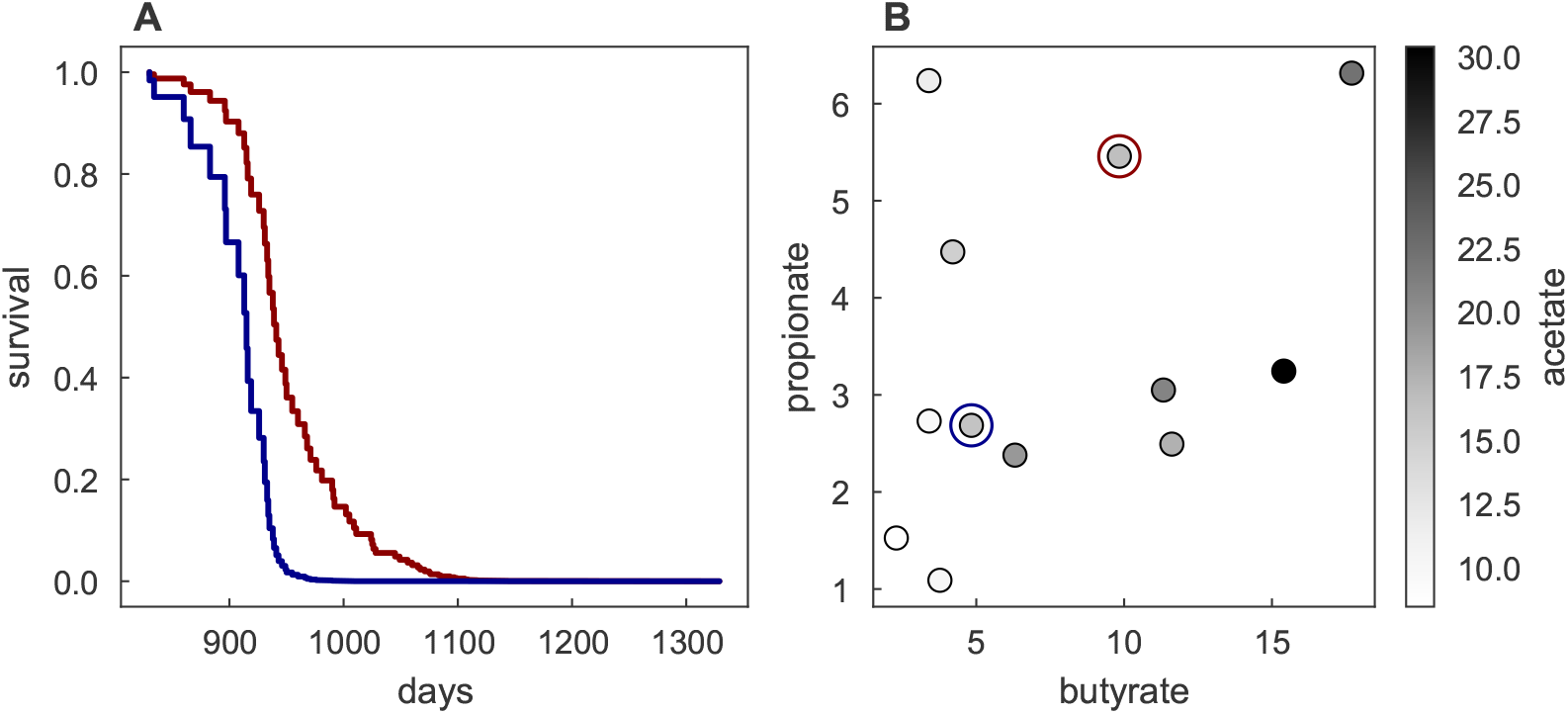
Effects of SCFAs on longevity. Predicted survival of mice exhibiting realistic variation in SCFA concentrations. (A) Expected survival curves for male, ACA treated mice at UM, conditional on being alive at 830 days of age, and (B) SCFA concentrations for that same set of mice. Two representative butyrate, propionate, and acetate concentrations were chosen to match the measured concentrations for a high (red) and low (blue) butyrate/propionate individual, both having similar acetate concentrations.

## List of Abbreviations

ACA: : Acarbose
SCFA: : Short-chain fatty acid
ITP: : Interventions Testing Program
TJL: : The Jackson Laboratory
UM: : The University of Michigan
UT: : The University of Texas Health Science Center at San Antonio
OTU: : Operational taxonomic unit
HPLC: : High-performance liquid chromatography
LASSO: : Least absolute shrinkage and selection operation
*S. alaskensis*: : *Sphingopyxis alaskensis*
FDR: : False-discovery rate
PBS: : Phosphate-buffered saline

## Ethics approval and consent to participate

All mice used in this study were maintained in specific-pathogen free conditions, and the protocols for husbandry and experimentation were approved by the Institutional Animal Care and Use Committees at each of the three institutions.

## Availability of data and materials

The sequence datasets generated and analyzed during the current study have been uploaded to the SRA database, accession SRP136736. Full-cohort survival data analyzed for portions of this study are available from the corresponding author on reasonable request. Code and metadata needed to reproduce the processing of raw data and downstream analyses is available on GitHub [44].

## Author’s contributions

DEH, RS, and RAM are principal ITP investigators at the three collaborating institutions and are responsible for design of the mouse experiment and supervision of technical personnel. ACE assembled and analyzed preliminary data. BJS collected the microbiome data and interpreted it along with TMS and RAM. BJS wrote a majority of the manuscript with contributions from TMS and RAM, who also helped with editing the manuscript. All authors read and approved the contents of the final article.

## Acknowledgments

We would like to thank Marian Schmidt, Nicole Koropatkin, Anna Seekatz, Nielson Baxter, Clive Waldron, and members of the Schmidt Lab for feedback. We appreciate the technical support at each of the study sites for maintaining the animals and collecting the samples.

